# *kcna1a* mutant zebrafish as a model of episodic ataxia type 1 and epilepsy

**DOI:** 10.1101/2022.09.28.509973

**Authors:** Deepika Dogra, Paola L. Meza-Santoscoy, Renata Rehak, Cristiane L R de la Hoz, Cezar Gavrilovici, Kingsley Ibhazehiebo, Jong M. Rho, Deborah M. Kurrasch

**Affiliations:** Department of Medical Genetics, University of Calgary, Calgary T2N 4N1, AB, Canada; Alberta Children’s Hospital Research Institute, University of Calgary, Calgary T2N 4N1, AB, Canada; Hotchkiss Brain Institute, University of Calgary, Calgary T2N 4N1, Alberta, Canada; Departments of Pediatrics, Clinical Neurosciences, Physiology & Pharmacology, Cumming School of Medicine, University of Calgary, Calgary T2N 4N1, Alberta, Canada; Departments of Neurosciences, Pediatrics, and Pharmacology, University of California San Diego, Rady Children’s Hospital San Diego, San Diego, CA 92037, USA

**Keywords:** episodic ataxia type 1, epilepsy, KCNA1, carbamazepine, zebrafish

## Abstract

**Objective:** *KCNA1* mutations are associated with a rare neurological movement disorder known as episodic ataxia type 1 (EA1), with epilepsy as a common comorbidity. Current medications only provide partial relief to ataxia and/or seizures, making new drugs needed. Here, we investigate the utility of zebrafish *kcna1a^−/−^* as a model of EA1 with epilepsy by characterizing its phenotype and comparing the efficacy of the first-line therapy carbamazepine in *kcna1a^−/−^* zebrafish to *Kcna1^−/−^* rodents.

**Methods:** We used CRISPR/Cas9 mutagenesis to introduce a mutation in the sixth segment of the zebrafish Kcna1 protein. Behavioral and electrophysiological assays were performed on *kcna1a^−/−^* larvae to assess ataxia- and epilepsy-related phenotypes. We also carried out real-time qPCRs to measure the transcript levels of brain hyperexcitability markers and bioenergetic profiling of *kcna1a^−/−^* larvae to evaluate their metabolic health. Carbamazepine efficacy was tested using behavioral assessments in *kcna1a^−/−^* zebrafish and seizure frequency in *Kcna1^−/−^* mice.

**Results:** *kcna1a^−/−^* zebrafish showed uncoordinated movements and locomotor deficits. The mutants also exhibited impaired startle responses when exposed to light-dark flashes and acoustic stimulation. Extracellular field recordings and upregulated *fosab* transcript levels showed hyperexcitability of the *kcna1a^−/−^* brain. Further, *vglut2a* and *gad1b* transcript levels were altered, indicative of neuronal excitatory/inhibitory imbalance in the *kcna1a^−/−^* brain. Metabolic health was also compromised in *kcna1a^−/−^* as seen by a significant reduction in measures of cellular respiration. Notably, carbamazepine reduced the impaired startle response in *kcna1a^−/−^* zebrafish but had no effect on the seizure frequency in *Kcna1^−/−^* mice, suggesting that this EA1 zebrafish model might better translate to human efficacy compared to rodents.

**Significance:** We conclude that zebrafish *kcna1a^−/−^* larvae show ataxia and epilepsy-related phenotypes and that they are responsive to carbamazepine treatment, consistent with EA1 patients. This study supports the notion that these zebrafish disease models can be useful for drug screening as well as studying the underlying disease biology.

**KEY POINTS:**

- Zebrafish *kcna1a^−/−^* larvae display dynamic behavioral changes, along with ataxia-like uncoordinated movements and brain hyperexcitability
- *kcna1a^−/−^* larvae have dysfunctional neuronal excitatory/inhibitory balance and perturbed metabolic health
- Similar to its effectiveness in patients, carbamazepine treatment improves behavioral deficits in *kcna1a^−/−^* larvae

## 1. INTRODUCTION

Mutations in the *KCNA1* gene, encoding voltage-gated potassium channel α-subunit Kv1.1, are associated with a rare neurological movement disorder known as episodic ataxia type 1 (EA1). The usual onset of EA1 occurs in childhood or early adolescence and patients often experience multiple attacks on daily basis (1). EA1 is characterized by myokymia and spastic contractions of skeletal muscles of head and limbs along with loss of motor coordination and balance. Some patients suffering from EA1 may also encounter focal-onset epilepsy, delayed motor development and cognitive disability. Episodes of discoordination in EA1 are usually triggered by stressors, including startle, emotional stress, exercise and temperature (2, 3).

KCNA1 is highly expressed in interneurons including basket cells of the cerebellum, where it forms inhibitory GABAergic synapses onto Purkinje cells. Specifically, KCNA1 plays an important role in the repolarization phase of presynaptic action potentials that provides the inhibitory inputs to Purkinje cells. Thus, *KCNA1* mutations result in the hyperexcitability of the presynaptic basket cells and the excessive release of GABA neurotransmitter, which can inhibit the generation of action potentials. As a consequence, the inhibitory output of the cerebellum may be markedly reduced, thus causing hyperexcitability and cerebellar symptoms observed in EA1 patients (4).

Pharmacological treatment of KCNA1-related EA1 is only symptomatic, with the carbonic anhydrase inhibitor acetazolamide providing some relief by reducing the frequency of ataxic episodes in some individuals. However, some long-term side effects of acetazolamide could include chronic metabolic acidosis, kidney stones, neuropsychiatric manifestations, fatigue and paresthesias (2, 5). The precise mechanism of action of acetazolamide in EA1 and other ion channelopathies is unexplored, although this drug is widely believed to alter the pH in the vicinity of the ion channel, causing hyperpolarization of the cell membrane and reducing neuronal hyperexcitability (6).

Phenytoin, a blocker of voltage-gated sodium channels is also effective in EA1 patients; however, its ability to induce permanent cerebellar dysfunction and atrophy has raised concerns against its long-term usage (5). Another sodium channel blocker, carbamazepine, can partially relieve ataxia and seizures in a patient subpopulation (7–9). Combined, current medications fail to provide effective treatment of the ataxia and seizures, making animal models necessary to provide further insights into the cellular and molecular mechanisms of the disorder as well as for drug screening to develop improved therapies.

Studies involving murine models of EA1 have revealed that the loss-of-function of KCNA1 causes an increase in GABA release, thereby inhibiting Purkinje cells in the cerebellum, ultimately leading to ataxia (10, 11). Furthermore, in support of an epileptogenic role for *KCNA1* mutations, *Kcna1^−/−^* mice exhibit epileptic phenotypes similar to patients, such as recurrent spontaneous seizures including myoclonic and generalized tonic-clonic seizures that begin 3-4 weeks postnatally (12, 13). *Kcna1^−/−^* mice also exhibit cardiorespiratory dysfunction and sudden unexpected death in epilepsy (14–17). *Kcna1^−/−^* mice have limited utility for drug screening owing to restricted animal numbers, longer reproductive periods, and the absence of high-throughput screening assays optimized for this model system. A more robust model of EA1 is needed to initiate large-scale drug screening required to discover better treatment options.

Zebrafish are commonly used to model human neurological disorders since zebrafish are vertebrates and display high molecular and cellular homology of the principal brain regions and associated networks comparable to humans. In addition, the behavior patterns of larval and adult zebrafish are well-established and can be easily analyzed, helpful in modeling various movement disorders. Several other advantages of using zebrafish include external fertilization, rapid development as well as the fact that they share >70% genes with humans and >80% of human disease-associated genes have a zebrafish counterpart (18). Thus, to explore the phenotypes related to EA1 and epilepsy due to loss of *KCNA1*, we generated a genetic mutant of one of the zebrafish paralogs *kcna1a*, expressed in reticulospinal neurons that regulate the animal’s startle response. We performed a comprehensive analysis of *kcna1a* homozygous mutants (hereafter addressed as *kcna1a^−/−^*) to obtain better insights into the behavioral, electrophysiological, molecular and metabolic consequences of KCNA1 dysfunction.

## 2. MATERIALS AND METHODS

### Zebrafish husbandry

All protocols and procedures were approved by the Health Science Animal Care Committee (protocol number AC18-0187) at the University of Calgary in compliance with the Guidelines of the Canadian Council of Animal Care. Adult wild-type zebrafish (TL strain) were maintained at 28°C in a 14-h light/10-h dark cycle under standard aquaculture conditions and fertilized eggs were collected via natural spawning. The animals were fed twice daily with Artemia. Zebrafish embryos and larvae were maintained in a non-CO_2_ incubator (VWR) at 28°C on the same light-dark cycle as the aquatic facility.

### Generation of zebrafish mutants

*kcna1a^ca201/ca201^* (*kcna1a^−/−^*) zebrafish were generated by using CRISPR/Cas9 mutagenesis. The identified founder carried a 6-nucleotide deletion within exon 2 (gRNA sequence: GGTTCCCTGTGCGCCATCGCTGG), producing a 2 amino acid deletion (Ile, Ala). The F1 heterozygous animals were outcrossed with wildtypes to raise F2 adults, which were used in the experiments. Genotyping was carried out by performing PCR using primers: *kcna1a*_F-GACCCTCAAAGCCAGTATGCG and *kcna1a*_R-GACTTGCTGACGGTTGAGGAG, followed by restriction digestion using BtgZI (NEB, R0703S). All three genotypes were distinguished by running the digested PCR product on a gel, based on different patterns and sizes of PCR product bands (wildtypes: 2 bands-214 and 191bp, heterozygotes: 3 bands-399, 214 and 191bp and homozygotes: 1 band-399bp). All the morphological analysis, including the eye and head size measurements were performed using ZEN software (ZEISS). Movies of freely swimming larvae were recorded using Stereo Discovery.V8 microscope (ZEISS).

### Gene expression analysis by in situ hybridization and real-time qPCR

DIG labeled probes were synthesized for *kcna1a* as previously described (19) using the following primers: *kcna1a*_insitu_F-CTCTGCCGTGCCGGGGCAT and *kcna1a*_insitu_R-CAATGTTCGGGTTGCTCACG. The embryos and larvae were fixed at requisite stage in 4% paraformaldehyde (Sigma-Aldrich) overnight at 4°C. The whole mount in situ hybridization protocol used was the same as described previously (19).

*fosab*, *vglut2a* and *gad1b* real-time qPCRs (qPCR) were performed on cDNA obtained from the heads of 3 dpf and 5 dpf larvae. *rpl13* was used as internal control. Following primers were used for qPCR: *rpl13*_qPCR_F-TAAGGACGGAGTGAACAACCA and *rpl13*_qPCR_R-CTTACGTCTGCGGATCTTTCTG; *fosab*_qPCR_F-TCGACGTGAACTCACCGATA *and fosab*_qPCR_R-CTTGCAGATGGGTTTGTGTG*; gad1b*_qPCR_F-AACTCAGGCGATTGTTGCA and *gad1b*_qPCR_R-TGAGGACATTTCCAGCCTTC; *vglut2a*_qPCR_F-CATCCTGTCTACAACTACGGTT and *vglut2a*_qPCR_R-CCAACACCAGAAATGAAATAGCCA.

### Behavioral assays

5 days post fertilization (dpf) zebrafish larvae maintained in 48-well plates were habituated for 20 minutes, under ambient light. This was followed by behavioral assessment to measure distance moved under various environmental conditions (100% light, light-dark flashes, acoustic startle) in Zebrabox (Viewpoint Life Sciences). Tracking of total distance moved as a measure of swimming behavior was analysed using Zebralab V3 software (Viewpoint Life Sciences). Distance travelled and time taken were retrieved from behavioral data to calculate speed.

### Drug treatment

Carbamazepine (Cayman Chemical) stock solution was prepared in DMSO. On the day of behavioral assays, the stock solution was diluted to a final concentration of 50 μM in embryo media. The zebrafish larvae were treated with 50 μM drug for 2 hours, followed by behavioral assessments. The final DMSO concentration was 0.5% and was used as vehicle control in behavioral assays.

For implant surgery and mouse video-EEG, carbamazepine was dissolved in DMSO to a final concentration of 40mg/kg body weight. Vehicle solution used was 10% DMSO.

### Metabolic measurements

Oxygen consumption rate measurements were performed using the XF24 Extracellular Flux Analyzer (Seahorse Biosciences). Single 3 dpf and 6 dpf zebrafish larvae were placed in 24 wells on an islet microplate and an islet plate capture screen was placed over the measurement area to maintain the larvae in place. Seven measurements were taken to establish basal rates, followed by treatment injections and 18 additional cycles (20). Rates were determined as the mean of two measurements, each lasting 2 min, with 3 min between each measurement (21). 3 independent assays were performed to establish metabolic measurements.

### Electrophysiological measurements

Electrophysiological recordings in zebrafish larvae were performed as previously described (22, 23). Briefly, 3 dpf zebrafish were paralysed using α-bungarotoxin (1 mg/ml, Tocris) and embedded in 1.2% low melting point agarose. The dorsal side of the zebrafish was exposed to the agarose gel surface and accessible for electrode placement. Larvae zebrafish were placed on an upright stage of a Zeiss Axioskop2 microscope, visualized using a 5X Zeiss N-Achroplan objective and perfused with embryo media. A glass microelectrode (3–8MΩ) filled with 2M NaCl was placed into the optic tectum of zebrafish, and recording was performed in current-clamp mode, low-pass filtered at 1 kHz, high-pass filtered at 0.1 Hz, using a digital gain of 10 (Multiclamp 700B amplifier, Digidata 1440A digitizer, Axon Instruments) and stored on a PC computer running pClamp software (Axon Instruments). Baseline recording was performed for 30 minutes.

### Implant surgery and mouse video-EEG

*Kcna1*-null mice were anaesthetized with 3% isoflurane, and the skull fixated with stereotaxic bars (KOPF Instruments) over heating pads (as approved in Animal protocol # AC21-0164). A 1.5 cm rostral-caudal incision was made to expose the skull surface, and connective tissues cleared. Three holes were gently drilled on each side of the skull with a 23mm gauge needle, avoiding suture lines. Mouse screws (0.1 inch) with wire leads were gently inserted midway into the skull, and a layer of dental acrylic applied over the screw heads and skull. A 6-pin mouse connector (Pinnacle Technology) was positioned over the screws, wires properly aligned, wrapped together, and folded backwards to avoid accidental contacts. A pocket for two EMG electrodes was made in the neck of the mouse by blunt dissection. Exposed wires were covered with a final layer of dental acrylic, and allowed to harden for 24 h before attaching the pin connector to the video EEG monitor (Pinnacle Technology). EEG signals were recorded simultaneously across the rostral, medial and caudal aspects of the neocortex using Sirenia Acquisition software (Pinnacle Technology). Mice were recorded 24 h per day for 1 day (baseline, postnatal day, P36-38), and received carbamazepine (intraperitoneal injection, 40 mg/kg body weight/day) for the next six consecutive days (P39–44). Frequency, duration, and time of seizure was analysed using Sirenia Seizure software (Pinnacle Technology).

### Statistical analysis

GraphPad Prism 9.4.1 was used to perform statistical analysis. Data are represented as mean ± s.e.m. P-values were calculated by unpaired t-test.

### Data availability

The authors declare that all data supporting the findings of this study are available within the article and its Supplementary Information files or from the corresponding author upon reasonable request.

## 3. RESULTS

### *kcna1a* mRNA expression in wild-type and morphological analyses of *kcna1a^−/−^* larvae

Zebrafish *kcna1a* is reported to be expressed in the Mauthner (M) cells that are a type of reticulospinal neurons (RSNs) in the hindbrain, contributing to neuronal firing and rapid startle behavior in response to external stimuli. *kcna1a* expression is also detected in two M cell homologs, i.e., MiD2cm and MiD3cm, involved in repetitive firing (24, 25). Here, by performing in situ hybridization, we further confirmed the *kcna1a* mRNA expression in the M cells in hindbrain at 2 dpf and 5 dpf, as well as in the RSNs in midbrain (nucleus of the medial longitudinal fasciculus nMLF) and neurons present in the spinal cord at 5 dpf (Figure 1A).

**Figure 1.**
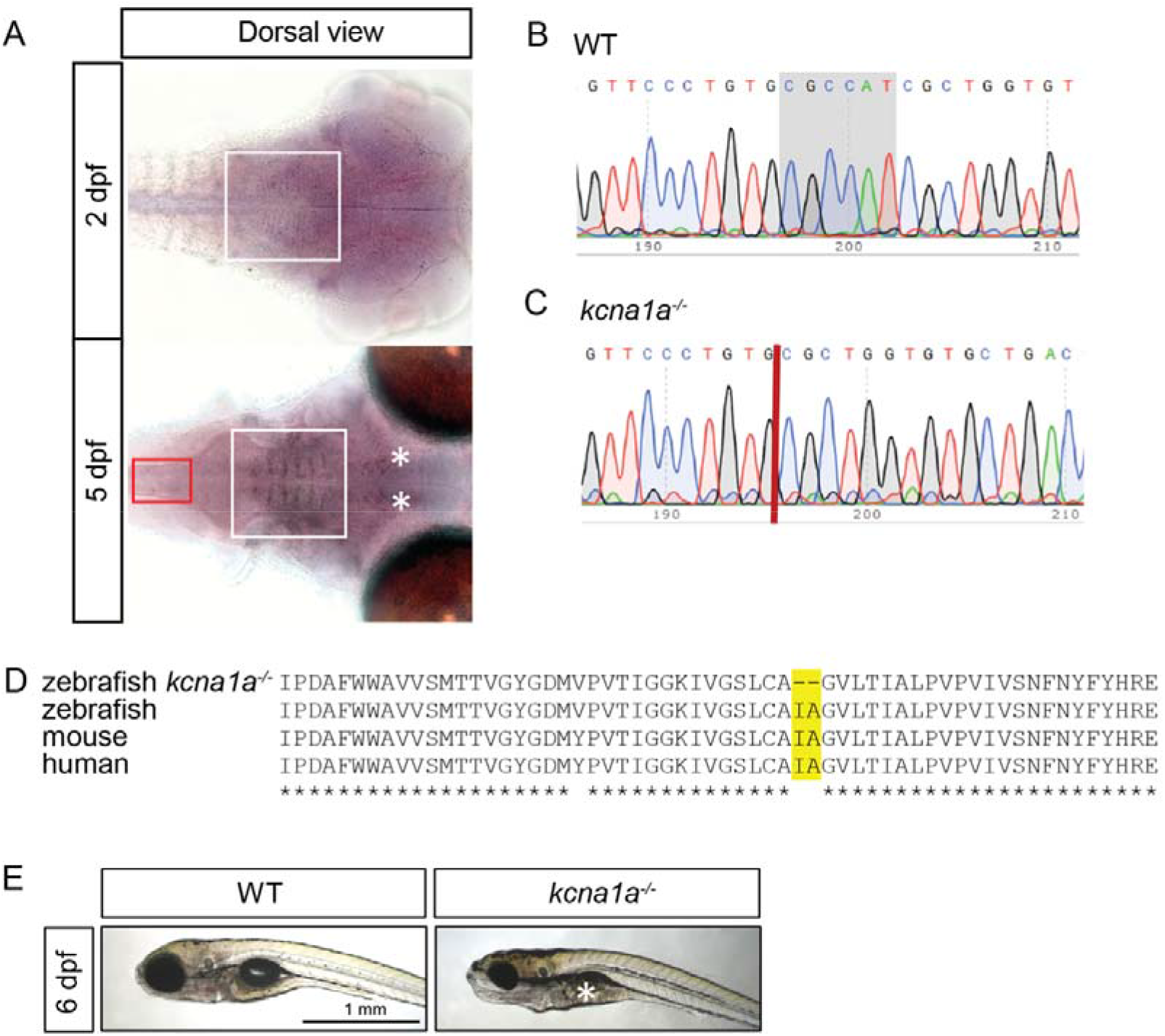
Expression analysis of *kcna1a* and generation of *kcna1a^−/−^* zebrafish model using CRISPR/Cas9 technique. (A) in situ hybridization for *kcna1a* expression at 2 dpf and 5 dpf. White boxes indicate RNA probe signal in M cells in hindbrain at 2 dpf and 5 dpf. White asterisks indicate RNA probe signal in nMLF in midbrain at 5 dpf. Red box indicates RNA probe signal in the neurons present in the spinal cord at 5 dpf. (B, C) Nucleotide sequences of *kcna1a* in zebrafish WT and *kcna1a^−/−^*. Grey highlighted region indicates the 6 nucleotides deleted due to mutation in *kcna1a^−/−^.* (D) Amino acid sequence of *kcna1a* in zebrafish *kcna1a^−/−^* and WT, and orthologues in mouse (Kcna1) and human (KCNA1). Yellow highlighted region indicates 2 amino acid deletion due to mutation in *kcna1a^−/−^*. (E) WT and *kcna1a^−/−^* larvae at 6 dpf. White asterisk indicating absence of swim bladder in *kcna1a^−/−^* larvae. Scale bars = 1 mm. dpf, days post fertilization; M cells, Mauthner cells; nMLF, nucleus of the medial longitudinal fasciculus: WT, wild-type.

Next, we injected a CRISPR targeting the second exon of *kcna1a* gene to generate a *kcna1a* mutant allele, *kcna1a^−/−^*. The mutation led to a 6-nucleotide deletion, causing a 2 amino acid deletion in the sixth transmembrane segment (S6) of the Kcna1a protein, which is conserved amongst humans, mice, and zebrafish (Figure 1B-D). Notably, mutations in the conserved Pro-Val-Pro motif in S6 are linked to epileptic encephalopathy (26, 27). Morphological analyses revealed no significant differences between *kcna1a^−/−^* larvae and wild-type siblings (WTs) at 3 dpf (Figure S1A). However, at 6 dpf, *kcna1a^−/−^* larvae exhibited a 13% reduction in eye diameter compared to WTs, with no significant difference in head size (Figure S1B,C). Interestingly, ~55% of 6 dpf *kcna1a^−/−^* larvae did not inflate their swim bladder, indicating developmental delay (Figure 1E, S1D). The homozygous mutants survive until 14 dpf (Figure S1E), whereas the heterozygotes live into adulthood and breed well (data not shown).

### *kcna1a^−/−^* larvae exhibit locomotor deficits and movement disorders

Zebrafish models of epilepsy exhibit behavioral comorbidities that model those observed in individuals with epilepsy (28). To ascertain the effect of *kcna1a* mutation on larval zebrafish behavior, we assessed larval swimming activity at 3 dpf and 5 dpf for 30 minutes in the light. The total distance travelled and speed of 3 dpf *kcna1a^−/−^* larvae were significantly higher compared to WTs (Figure 2A, B). The locomotor plots of 3 dpf *kcna1a^−/−^* larvae further showed hyperactivity compared to WTs that swim little during this stage (Figure 2C). Conversely, when assessed at 5 dpf, *kcna1a^−/−^* larvae swam significantly shorter distances and with decreased speed compared to WTs (Figure 2D, E). The locomotor plots further depict that 5 dpf *kcna1a^−/−^* larvae were less active and displayed an uncoordinated movement pattern, a sign of ataxia (Figure 2F). We further analyzed the pattern of swimming trajectory in 5 dpf mutants by recording movies and confirmed that their swimming pattern was abnormally distinct from WTs (Movie S1,S2). Taken together, these results illustrate the dynamic changes in swimming behaviors of *kcna1a^−/−^* larvae during different stages of development and their ataxia-like features that appear at a later age.

**Figure 2.**
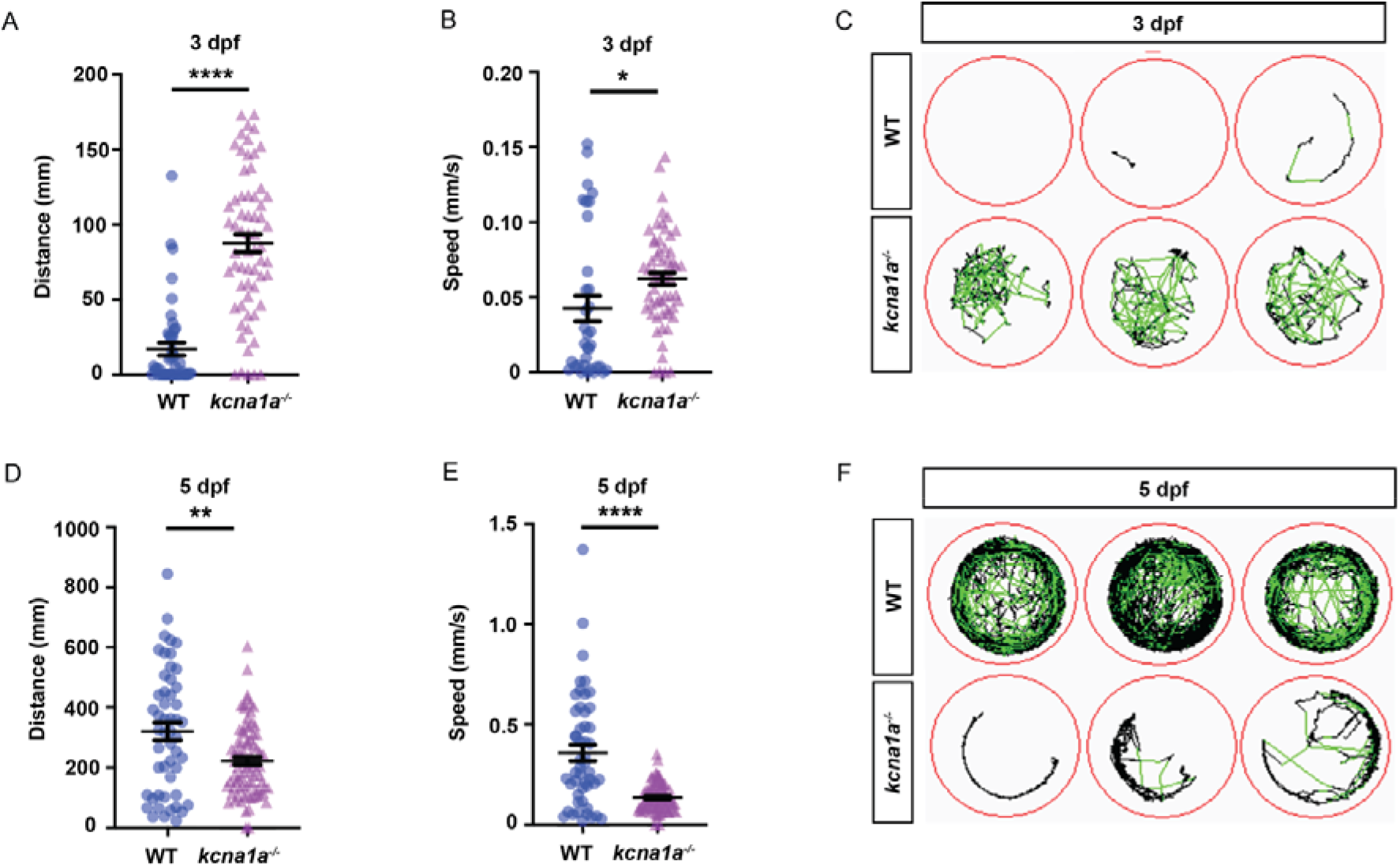
Spontaneous behavior analysis of *kcna1a^−/−^* zebrafish. (A) Quantification of distance travelled at 3 dpf tracked in 100% light for 30 minutes. *kcna1a^−/−^* move higher distances compared to WT. WT, n=44; *kcna1a^−/−^,* n=63. (B) Quantification of speed at 3 dpf tracked in 100% light for 30 minutes. Speed of *kcna1a^−/−^* is significantly higher than WT. WT, n=34; *kcna1a^−/−^,* n=60. (C) Locomotor plots at 3 dpf, showing that *kcna1a^−/−^* are more active than WTs. WT, n=20; *kcna1a^−/−^,* n=20. (D) Quantification of distance travelled at 5 dpf tracked in 100% light for 30 minutes. Distance travelled by *kcna1a^−/−^* is significantly reduced compared to WT. WT, n=51; *kcna1a^−/−^,* n=84. (E) Quantification of speed at 5 dpf tracked in 100% light for 30 minutes. Speed of *kcna1a^−/−^* is significantly reduced compared to WT. WT, n=50; *kcna1a^−/−^,* n=84. (F) Locomotor plots at 5 dpf, showing that *kcna1a^−/−^* are moving less and have abnormal swimming patterns. WT, n=20; *kcna1a^−/−^,* n=24. Data are mean ± s.e.m., *P ≤ 0.05, **P ≤ 0.01, ****P ≤ 0.0001-Unpaired t-test.

### *kcna1a^−/−^* larvae exhibit impaired startle response that is rescued by carbamazepine treatment

Since convulsions in the EA1 patients can be triggered by extrinsic stimuli, we assessed the response of *kcna1a^−/−^* larvae to different environmental triggers. We tested an acoustic startle protocol on 5 dpf larvae for 10 minutes by introducing vibration pulses of 440Hz in light (Figure 3A) and calculated the total distance travelled at the end of every 1 minute. The *kcna1a^−/−^* larvae moved significantly longer distances during the second and third pulses compared to WTs, consistent with an abnormal response to acoustic startle (Figure 3B) and a recent study (29). We also performed a protocol of light-dark flashes on 5 dpf larvae for a duration of 90 seconds (Figure 3C) and measured the total distance travelled at the end of every 3 seconds. The *kcna1a^−/−^* larvae showed an exaggerated response to light-dark flashes and traveled longer distances compared to WTs (Figure 3D).

**Figure 3.**
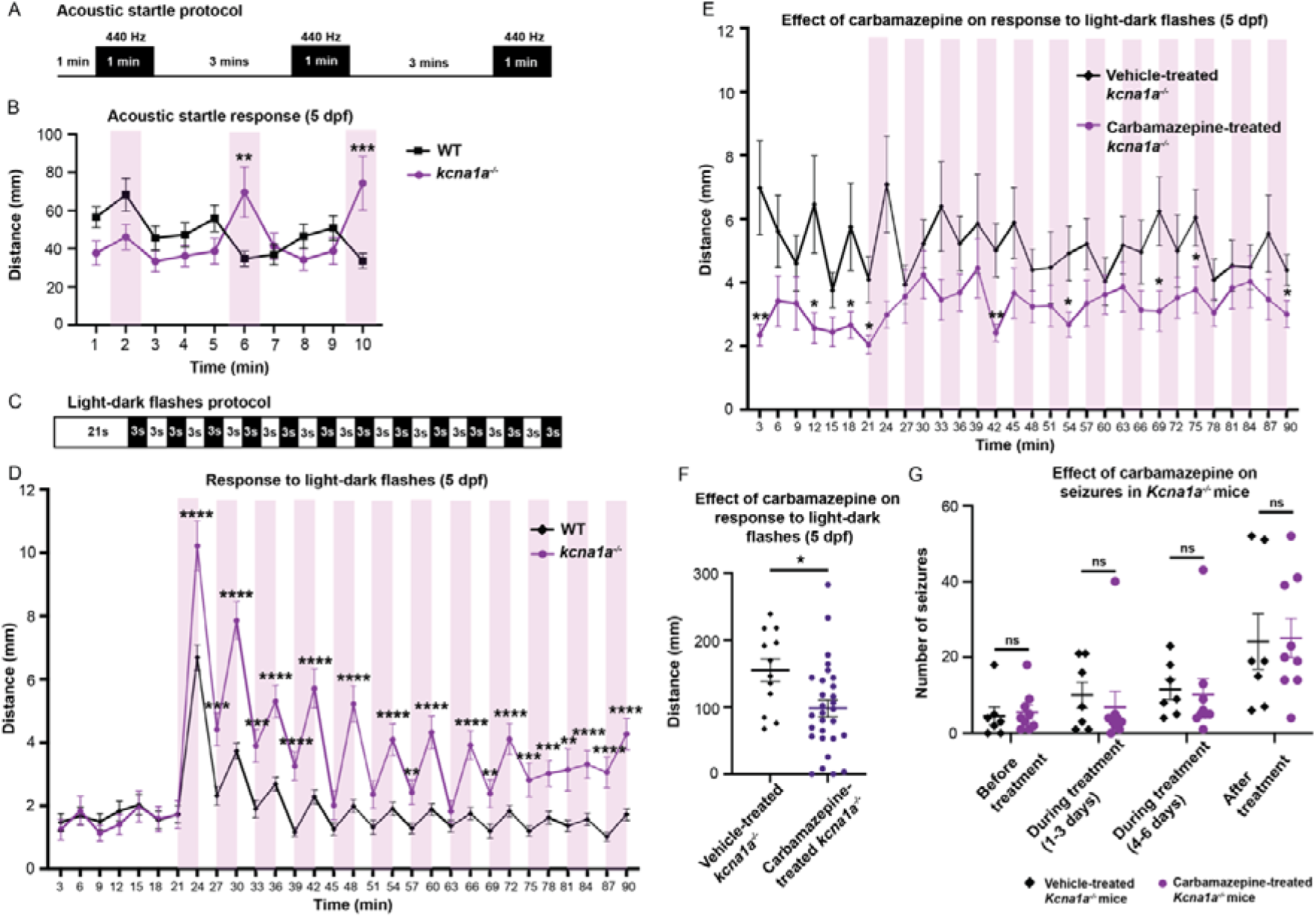
Analysis of startle response in *kcna1a^−/−^* zebrafish and effect of carbamazepine treatment on *kcna1a^−/−^* zebrafish and *Kcna1a^−/−^* mice. (A) Schematic representation of acoustic startle protocol. (B) Quantification of distance travelled every 1 minute in acoustic startle at 5 dpf. *kcna1a^−/−^* are more responsive to startle with significantly higher distance travelled in second and third vibration pulse compared to WT. WT, n=113; *kcna1a^−/−^,* n=52. (C) Schematic representation of light-dark flashes protocol. (D) Quantification of distance travelled every 3 seconds in light-dark flashes at 5 dpf. *kcna1a^−/−^* show higher sensitivity to startle with more distance travelled compared to WT. WT, n=108; *kcna1a^−/−^,* n=63. (E, F) Quantification of distance travelled every 3 seconds (E) and in 90 seconds (F) in light-dark flashes after treatment with carbamazepine at 5 dpf. Carbamazepine treatment rescues impaired startle response in *kcna1a^−/−^*. Vehicle-treated *kcna1a^−/−^*, n=12; carbamazepine-treated *kcna1a^−/−^,* n=29. (G) Quantification of number of seizures encountered by vehicle- and carbamazepine-treated P36-38 *Kcna1a^−/−^* mice at different time points. Carbamazepine treatment does not reduce the frequency of seizures in *Kcna1a^−/−^* mice. Vehicle-treated *Kcna1a^−/−^*, n=7; carbamazepine-treated *Kcna1a^−/−^,* n=9. Data are mean ± s.e.m., ns: no significant changes observed, *P ≤ 0.05, **P ≤ 0.01, ***P ≤ 0.001, ****P ≤ 0.0001-Unpaired t-test.

Next, we examined whether the impaired startle behavior of our *kcna1a^−/−^* zebrafish larvae was improved by treatment with carbamazepine, a sodium channel blocker that can reduce the neuronal hyperexcitability in patients with KCNA1-related epilepsy (7). We treated 5 dpf *kcna1a^−/−^* larvae with carbamazepine (50 μM) and DMSO (vehicle-control; 0.5%) for 2 hours and performed the light-dark flashes protocol (as described above) to assess the distance travelled. Notably, this dose was selected by titrating carbamazepine to identify the concentration that did not significantly affect the swimming activity of WTs (data not shown). Carbamazepine treatment rescued the impaired startle response in *kcna1a^−/−^* larvae since the distance travelled every 3 seconds was significantly lower at several time-points and the total distance swam during the complete protocol of 90 seconds was reduced by ~37% (Figure 3E,F).

Mouse models are traditionally favored over zebrafish for translational studies and thus we wanted to compare the efficacy of carbamazepine in *kcna1^−/−^* zebrafish to its effects in *Kcna1^−/−^* mice that display spontaneous recurrent seizures. Previous studies using *Kcna1* knockout mice have shown that carbamazepine had partial efficacy in seizure reduction but did not eliminate severe seizures (30, 31). We treated epileptic P36-38 *Kcna1^−/−^* mice with carbamazepine (40 mg/kg) or DMSO (vehicle-control; 10%) and quantified the number of seizures per day with video-EEG recordings across an early-, late-, and post-treatment window (Figure 3G). Carbamazepine treatment had no effect on seizure frequency in *Kcna1^−/−^* mice, indicating that, at least for this drug, the zebrafish larvae response is more in alignment with *KCNA1* patients than mice.

### *kcna1a^−/−^* larvae exhibit brain hyperexcitability and excitatory/inhibitory neuron imbalance

Pharmacological and genetic epilepsy models of zebrafish display brain hyperexcitability in electrophysiological recordings as reported in several studies (22, 23), similar to mouse models and humans with epilepsy. Thus, we also wanted to determine whether a *kcna1a* mutation causes brain hyperactivity, which serves as a proxy for an epileptic state. We measured extracellular field potentials from the optic tectum of agarose-immobilized 3 dpf WTs and *kcna1a^−/−^* larvae brains (Figure 4A). WTs showed no evidence of abnormal electrical activity; however, extracellular field recordings in ~60% of *kcna1a^−/−^* larvae revealed repetitive high-frequency, large-amplitude spikes, consistent with a spontaneous epileptic phenotype (Figure 4B,C,D). Since seizures induced in a rodent model can lead to an upregulated expression of immediate early genes (IEGs) in brain regions where the seizures develop (32), we measured the expression of an IEG, *fosab,* at 3 dpf and 5 dpf by performing qPCR on cDNA obtained from larvae heads. Compared to WTs, *kcna1a^−/−^* larvae brains showed a significant upregulation in the *fosab* transcript levels at 3 dpf and 5 dpf, another confirmation of brain hyperexcitability (Figure 4E). Recent studies on zebrafish epilepsy models have also revealed that the neuronal excitatory/inhibitory (E/I) balance is disturbed in epileptic brain (33, 34). Thus, we further quantified excitatory glutamatergic (*vglut2a*) and inhibitory GABAergic (*gad1b*) neuronal marker expression in the larval brains by performing qPCR at 3 dpf and 5 dpf. The expression of *vglut2a* was significantly upregulated; however, *gad1b* transcript levels were significantly reduced in *kcna1a^−/−^* larvae compared to WTs at 3 dpf and 5 dpf, indicative of excitatory/inhibitory (E/I) imbalance in neurons due to mutation in *kcna1a* (Figure 4F,G).

**Figure 4.**
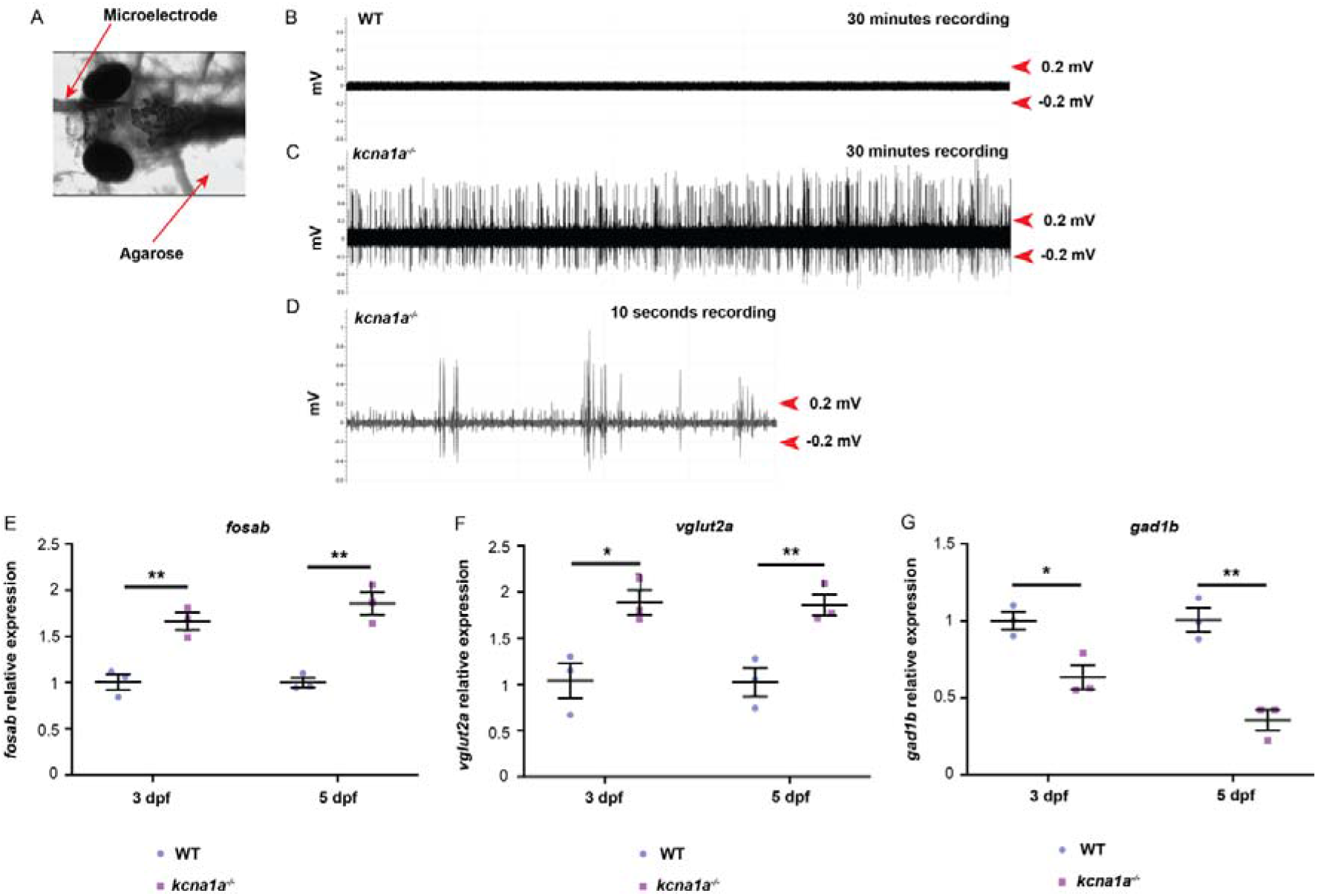
Brain hyperexcitability and E/I imbalance analysis in *kcna1a^−/−^* zebrafish. (A) Image of agarose-embedded zebrafish with glass microelectrode placed in the optic tectum. (B, C, D) Representative extracellular field recordings obtained from the optic tectum of 3 dpf WT (B) and *kcna1a^−/−^* zebrafish larvae (C, D-higher magnification of C). The high frequency large amplitude epileptiform-like discharges seen in the *kcna1a^−/−^* (that are absent in the WT) are indicative of increased network hyperexcitability. WT, n=4; *kcna1a^−/−^,* n=7. (E, F, G) qPCR analysis for relative *fosab, vglut2a* and *gad1b* mRNA expression in 3 dpf and 5 dpf *kcna1a^−/−^* larvae compared to WT. WT and *kcna1a^−/−^*, n = 3 × 10 larvae assessed as three biological and two technical replicates each. *fosab* is upregulated in *kcna1a^−/−^* indicating brain hyperexcitability. *vglut2a* is upregulated and *gad1b* is downregulated in *kcna1a^−/−^* indicating dysfunctional E/I balance. Data are mean ± s.e.m., ns: no significant changes observed, *P ≤ 0.05, **P ≤ 0.01-Unpaired t-test.

### *kcna1a^−/−^* larvae experience compromised mitochondrial health

Studies on cortical cell cultures generated from rodent brains demonstrate that prolonged seizure activity negatively affects mitochondrial bioenergetics and induces cell death (35). Our group and others also show that mitochondrial bioenergetics is altered in various epileptic models of zebrafish (23, 36, 37). In this study, to assess whether mitochondrial bioenergetics is dysregulated in our *kcna1a^−/−^* larvae, we used the Seahorse XF Flux Bioanalyzer to measure metabolic changes in zebrafish larvae at 3 dpf and 6 dpf (Figure 5A). We found that the basal respiration was significantly decreased in *kcna1a^−/−^* larvae at 3 dpf and 6 dpf compared to WTs (Figure 5B). Moreover, a significant reduction in mitochondrial respiration at 6 dpf (unchanged at 3 dpf) and a significant reduction in non-mitochondrial respiration at 3 dpf (unchanged at 6 dpf) in *kcna1a^−/−^* larvae compared to WTs (Figure 5C,D) was observed. These findings collectively indicate that *kcna1a^−/−^* larvae experience decreased energy utilization, which is consistent with brain PET scan findings in a patient with EA1 due to *KCNA1* mutation (38).

**Figure 5.**
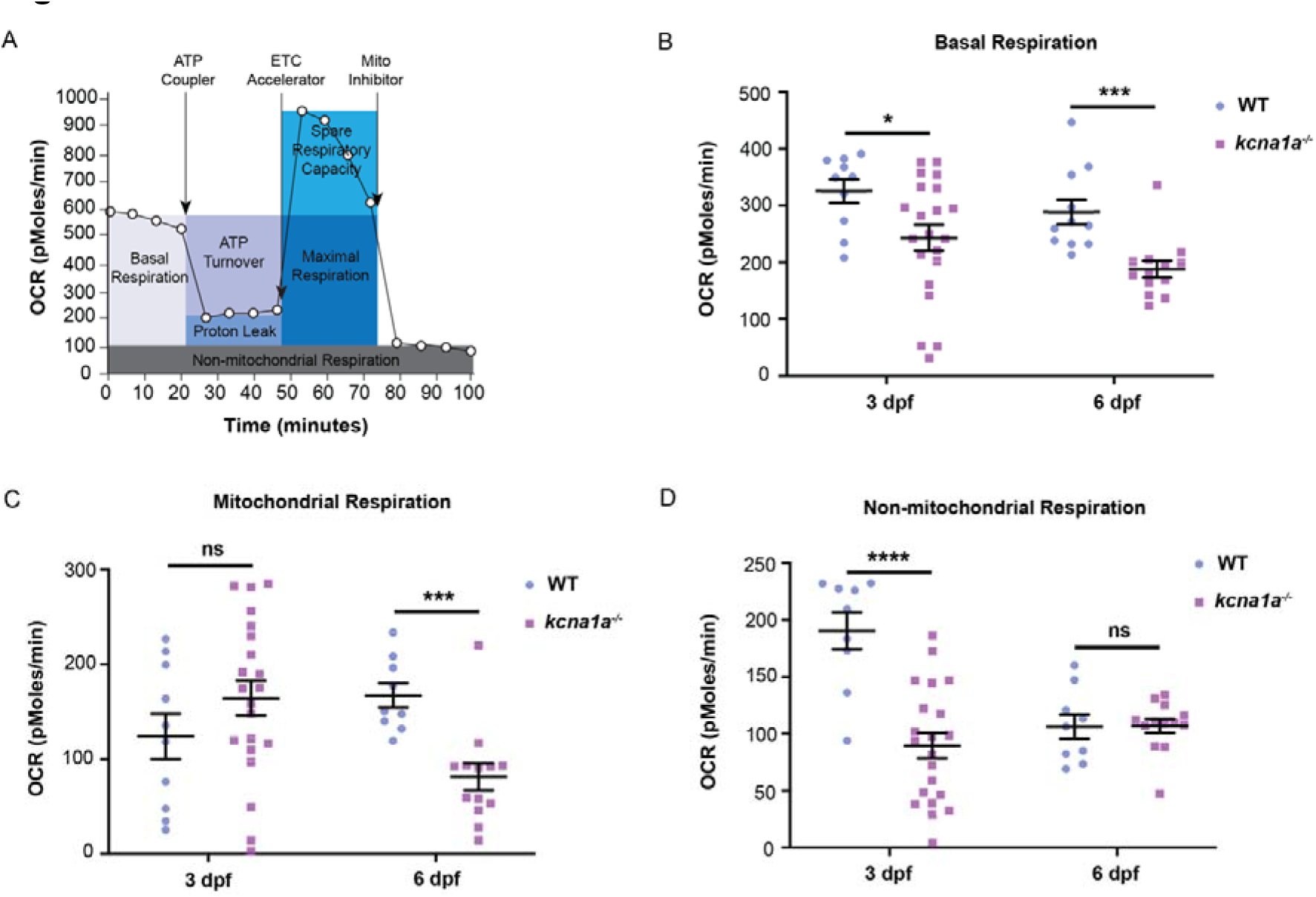
Metabolic characterization of *kcna1a^−/−^* zebrafish. (A) Schematic representation of how the Seahorse bioanalyser displays mitochondria bioenergetics as regulated by pharmacological inhibitors. (B) Quantification of basal respiration. *kcna1a^−/−^* exhibit a significant reduction in basal respiration compared to WT at 3 dpf and 6 dpf. WT, n=10; *kcna1a^−/−^,* n=21 at 3 dpf. WT, n=11; *kcna1a^−/−^,* n=13 at 6 dpf. (C) Quantification of mitochondrial respiration. *kcna1a^−/−^* exhibit a significant reduction in mitochondrial respiration compared to WT at 6 dpf. WT, n=10; *kcna1a^−/−^,* n=21 at 3 dpf. WT, n=9; *kcna1a^−/−^,* n=13 at 6 dpf. (D) Quantification of non-mitochondrial respiration. *kcna1a^−/−^* exhibit a significant reduction in non-mitochondrial respiration compared to WT at 3 dpf. WT, n=19; *kcna1a^−/−^,* n=21 at 3 dpf. WT, n=9; *kcna1a^−/−^,* n=13 at 6 dpf. Data are mean ± s.e.m., ns: no significant changes observed, *P ≤ 0.05, ***P ≤ 0.001, ****P ≤ 0.0001-Unpaired t-test.

## 4. DISCUSSION

Currently, there are no standard treatment options for EA1 patients and thus efforts should be made to identify new and better medications. In this study, we generated a *kcna1a* loss-of-function zebrafish model of EA1 and epilepsy. Our zebrafish mutants are also responsive to the existing first-line treatment of carbamazepine. We propose that our current zebrafish model can be further incorporated for understanding the biology of this disorder as well as drug screening to identify more effective therapies.

To date, *KCNA1* is the only gene associated with EA1, with more than 40 *KCNA1* loss-of-function missense mutations reported (5, 39). Although a range of symptoms are caused by variants in *KCNA1*, over 50% of all variants result in a diagnosis of EA1 and the remaining 50% associated with additional comorbidities, including EA1 with epilepsy, epilepsy alone, myokymia, hyperthermia and hypomagnesemia (26, 40, 41). Predictably, patients with distinct *KCNA1* variants have differential responses to drug treatments, confirmed by a recent report that compiled the available clinical findings of 15 patients with 36 treatment efforts using 12 different drugs (9). The authors found that two sodium channel blockers were promising, with carbamazepine showing the most therapeutic benefits and phenytoin leading to substantial clinical improvement. Acetazolamide displayed a mixed response, with ~45% patients showing positive effects but the majority without any improvement. The field currently lacks a model system that phenocopies the patient and that can be robustly employed in large-scale screening for new drugs. We propose that our *kcna1a^−/−^* zebrafish are a useful model to study the biology and identification of treatments for individuals that harbor a *KCNA1* mutation.

We previously used *kcna1a* zebrafish morphants to test our anti-seizure drug screening platform. *kcna1a* morphants demonstrate hyperactivity when tracked in dark, as well as high frequency, high amplitude spikes in extracellular field recordings. Amongst several compounds screened, Vorinostat, a histone deacetylase inhibitor was selected as a candidate drug for treating epilepsy as it significantly restores dysregulated bioenergetics parameters and reduces brain hyperexcitability in *kcna1a* morphants (23). Our present study describes a new zebrafish *kcna1a^−/−^* model that recapitulates many aspects of EA1 and epilepsy observed in patients suffering from *KCNA1* mutations. Our zebrafish *kcna1a^−/−^* larvae show dynamic behavioral deficits, with enhanced swimming activity in early stages and reduced swimming activity with uncoordinated movement at later stages of development (indicative of ataxia). Similar abnormal swimming patterns have been reported in another zebrafish model of ataxia (42). Further, the presence of hyperexcitability and bursting in extracellular field recordings, along with upregulated mRNA expression of *fosab* and perturbed E/I balance, is consistent with an epilepsy phenotype in these mutants. The early death of our *kcna1a^−/−^* zebrafish correlates with the phenomenon of sudden unexpected death in epilepsy (SUDEP), which also occurs in neuron-specific *Kcna1* knockout mice (43). The abnormal startle behavior exhibited by *kcna1a^−/−^* larvae mimics the spasms triggered by EA1 patients in response to external stimuli. Similar deficits in acoustic startle response and habituation learning have been reported in other *kcna1a^−/−^* zebrafish models (29, 44). Moreover, carbamazepine treatment in *kcna1a^−/−^* zebrafish rescues this impaired startle response, whereas treatment in *Kcna1^−/−^* mice had no effect on seizure frequency, indicating that *kcna1a^−/−^* zebrafish might better phenocopy EA1 patients, at least in their responsiveness to this therapeutic.

Our bioenergetics analysis indicates that the *kcna1a^−/−^* zebrafish exhibit a significant reduction in neurometabolism. Interestingly, we previously observed opposite OCR effects in the *kcna1a* morpholino-injected larvae, which show an increase in basal respiration, total mitochondrial respiration and ATP-linked respiration (23). These differences could be possibly due to variability in the epilepsy models used, i.e., acute seizure-inducing *kcna1a* morphants and versus chronic seizure-inducing *kcna1a* genetic mutants. Such differences in patterns of metabolic deficits have also been reported previously using different zebrafish models of epilepsy (37). The underlying reason(s) for these differences remain unclear.

Importantly, our *kcna1a^−/−^* zebrafish holds a promising role for drug screening as well as extensive transcriptomic/proteomics profiling for gaining new insights into the molecular mechanisms behind this disorder. We are currently employing these genetic mutants in screening the combinations of approved anti-seizure medications. Several research groups show that drugs that are functional in humans have a same target in zebrafish, including targeted cancer therapies like MEK inhibitors (45) and serotonin modulators (46). More than two decades of zebrafish research in drug screening suggests that most of the molecules that are active in zebrafish are also active in humans and rodents (47). Thus, zebrafish hold the potential to be extensively utilized for drug repurposing and precision medicine-based drug discovery.

The field can further benefit by generating new *kcna1a* loss-of-function zebrafish models recapitulating patients’ mutations to unravel the differentially regulated signaling pathways as well as to compare their responsiveness to different drug treatments. Precision gene editing can help generate patient-specific models in zebrafish and these models could recapitulate the etiology found in the patients (48). We must, however, remember the shortcomings of using zebrafish as a model system in isolation, including the absence of corticospinal and rubrospinal tracts in the zebrafish CNS (49) and gene duplication leading to genetic redundancy that could complicate model development (18). Therefore, efforts should be made to incorporate more model systems like rodents and/or patient-specific brain organoids (50) to further validate top drug candidates from a zebrafish-based drug screening before they reach clinical trials.

## 5. CONCLUSIONS

We conclude that our *kcna1a^−/−^* larvae show several phenotypes that model the patient symptoms including ataxia, epilepsy and locomotion behavioral deficits. Further, the fact that these zebrafish mutants are responsive to carbamazepine treatment similar to EA1 patients shows a translation of this EA1 model. Our study supports the idea that zebrafish epilepsy disease models are valuable as a drug screening platform, suitable for both testing new therapeutical approaches and dissecting the underlying etiology of this disorder.

## AUTHOR CONTRIBUTIONS

D.D. designed the study, performed the experiments, analyzed the data and wrote the manuscript; P.L.M.S. performed some of the behavioral assays and in situ hybridization experiments; R.R. generated the zebrafish mutants and performed some of the mice experiments; C.L.R.D.L.H. performed some of the mice experiments, K.I. performed bioenergetics experiments; C.G. and J.M.R. performed and planned electrophysiological experiments, respectively; D.M.K. designed the study, analyzed the data, wrote the manuscript and supervised the work. All the authors commented on the manuscript.

## ACKNOWLEDGMENTS

We would like to thank Dr. Timothy Simeone (Creighton University) for providing *Kcna1^+/−^* breeders. We further thank Arthur Omorogiuwa for assistance with zebrafish husbandry and and Natasha Klenin for assistance with zebrafish genotyping. We also thank Elizabeth Hughes for assistance with mouse husbandry and genotyping and Younghee Ahn for training and support on the Seahorse XF Bioanalyzer. Finally, we thank Alicia Vandenbrink for administrative support.

## SUPPLEMENTARY MATERIAL

Supplementary material is available online.

## REFERENCES

1. Rajakulendran S, Schorge S, Kullmann DM, Hanna MG. Episodic ataxia type 1: a neuronal potassium channelopathy. Neurotherapeutics. 2007;4(2):258–66.

2. Graves TD, Cha YH, Hahn AF, Barohn R, Salajegheh MK, Griggs RC, et al. Episodic ataxia type 1: clinical characterization, quality of life and genotype-phenotype correlation. Brain. 2014;137(Pt 4):1009–18.

3. Imbrici P, Altamura C, Gualandi F, Mangiatordi GF, Neri M, De Maria G, et al. A novel KCNA1 mutation in a patient with paroxysmal ataxia, myokymia, painful contractures and metabolic dysfunctions. Mol Cell Neurosci. 2017;83:6–12.

4. Choi KD, Choi JH. Episodic Ataxias: Clinical and Genetic Features. J Mov Disord. 2016;9(3):129–35.

5. D'Adamo MC, Hasan S, Guglielmi L, Servettini I, Cenciarini M, Catacuzzeno L, et al. New insights into the pathogenesis and therapeutics of episodic ataxia type 1. Front Cell Neurosci. 2015;9:317.

6. Brunt ER, van Weerden TW. Familial paroxysmal kinesigenic ataxia and continuous myokymia. Brain. 1990;113 (Pt 5):1361–82.

7. Miceli F, Guerrini R, Nappi M, Soldovieri MV, Cellini E, Gurnett CA, et al. Distinct epilepsy phenotypes and response to drugs in KCNA1 gain-and loss-of function variants. Epilepsia. 2022;63(1):e7–e14.

8. Maggi L, Bonanno S, Altamura C, Desaphy JF. Ion Channel Gene Mutations Causing Skeletal Muscle Disorders: Pathomechanisms and Opportunities for Therapy. Cells. 2021;10(6).

9. Lauxmann S, Sonnenberg L, Koch NA, Bosselmann C, Winter N, Schwarz N, et al. Therapeutic Potential of Sodium Channel Blockers as a Targeted Therapy Approach in KCNA1-Associated Episodic Ataxia and a Comprehensive Review of the Literature. Front Neurol. 2021;12:703970.

10. Herson PS, Virk M, Rustay NR, Bond CT, Crabbe JC, Adelman JP, et al. A mouse model of episodic ataxia type-1. Nat Neurosci. 2003;6(4):378–83.

11. Begum R, Bakiri Y, Volynski KE, Kullmann DM. Action potential broadening in a presynaptic channelopathy. Nat Commun. 2016;7:12102.

12. Smart SL, Lopantsev V, Zhang CL, Robbins CA, Wang H, Chiu SY, et al. Deletion of the K(V)1.1 potassium channel causes epilepsy in mice. Neuron. 1998;20(4):809–19.

13. Rho JM, Szot P, Tempel BL, Schwartzkroin PA. Developmental seizure susceptibility of kv1.1 potassium channel knockout mice. Dev Neurosci. 1999;21(3–5):320–7.

14. Glasscock E, Yoo JW, Chen TT, Klassen TL, Noebels JL. Kv1.1 potassium channel deficiency reveals brain-driven cardiac dysfunction as a candidate mechanism for sudden unexplained death in epilepsy. J Neurosci. 2010;30(15):5167–75.

15. Aiba I, Noebels JL. Spreading depolarization in the brainstem mediates sudden cardiorespiratory arrest in mouse SUDEP models. Sci Transl Med. 2015;7(282):282ra46.

16. Simeone KA, Hallgren J, Bockman CS, Aggarwal A, Kansal V, Netzel L, et al. Respiratory dysfunction progresses with age in Kcna1-null mice, a model of sudden unexpected death in epilepsy. Epilepsia. 2018;59(2):345–57.

17. Dhaibar H, Gautier NM, Chernyshev OY, Dominic P, Glasscock E. Cardiorespiratory profiling reveals primary breathing dysfunction in Kcna1-null mice: Implications for sudden unexpected death in epilepsy. Neurobiol Dis. 2019;127:502–11.

18. Howe K, Clark MD, Torroja CF, Torrance J, Berthelot C, Muffato M, et al. The zebrafish reference genome sequence and its relationship to the human genome. Nature. 2013;496(7446):498–503.

19. Thisse C, Thisse B. High-resolution in situ hybridization to whole-mount zebrafish embryos. Nat Protoc. 2008;3(1):59–69.

20. Stackley KD, Beeson CC, Rahn JJ, Chan SS. Bioenergetic profiling of zebrafish embryonic development. PLoS One. 2011;6(9):e25652.

21. Richon VM. Targeting histone deacetylases: development of vorinostat for the treatment of cancer. Epigenomics. 2010;2(3):457–65.

22. Baraban SC, Dinday MT, Castro PA, Chege S, Guyenet S, Taylor MR. A large-scale mutagenesis screen to identify seizure-resistant zebrafish. Epilepsia. 2007;48(6):1151–7.

23. Ibhazehiebo K, Gavrilovici C, de la Hoz CL, Ma SC, Rehak R, Kaushik G, et al. A novel metabolism-based phenotypic drug discovery platform in zebrafish uncovers HDACs 1 and 3 as a potential combined anti-seizure drug target. Brain. 2018;141(3):744–61.

24. Brewster DL, Ali DW. Expression of the voltage-gated potassium channel subunit Kv1.1 in embryonic zebrafish Mauthner cells. Neurosci Lett. 2013;539:54–9.

25. Watanabe T, Shimazaki T, Mishiro A, Suzuki T, Hirata H, Tanimoto M, et al. Coexpression of auxiliary Kvbeta2 subunits with Kv1.1 channels is required for developmental acquisition of unique firing properties of zebrafish Mauthner cells. J Neurophysiol. 2014;111(6):1153–64.

26. Rogers A, Golumbek P, Cellini E, Doccini V, Guerrini R, Wallgren-Pettersson C, et al. De novo KCNA1 variants in the PVP motif cause infantile epileptic encephalopathy and cognitive impairment similar to recurrent KCNA2 variants. Am J Med Genet A. 2018;176(8):1748–52.

27. Russo A, Gobbi G, Pini A, Moller RS, Rubboli G. Encephalopathy related to status epilepticus during sleep due to a de novo KCNA1 variant in the Kv-specific Pro-Val-Pro motif: phenotypic description and remarkable electroclinical response to ACTH. Epileptic Disord. 2020;22(6):802–6.

28. Grone BP, Qu T, Baraban SC. Behavioral Comorbidities and Drug Treatments in a Zebrafish scn1lab Model of Dravet Syndrome. eNeuro. 2017;4(4).

29. Meserve JH, Nelson JC, Marsden KC, Hsu J, Echeverry FA, Jain RA, et al. A forward genetic screen identifies Dolk as a regulator of startle magnitude through the potassium channel subunit Kv1.1. PLoS Genet. 2021;17(6):e1008943.

30. Lavebratt C, Trifunovski A, Persson AS, Wang FH, Klason T, Ohman I, et al. Carbamazepine protects against megencephaly and abnormal expression of BDNF and Nogo signaling components in the mceph/mceph mouse. Neurobiol Dis. 2006;24(2):374–83.

31. Deodhar M, Matthews SA, Thomas B, Adamian L, Mattes S, Wells T, et al. Pharmacoresponsiveness of spontaneous recurrent seizures and the comorbid sleep disorder of epileptic Kcna1-null mice. Eur J Pharmacol. 2021;913:174656.

32. Morgan JI, Cohen DR, Hempstead JL, Curran T. Mapping patterns of c-fos expression in the central nervous system after seizure. Science. 1987;237(4811):192–7.

33. Brenet A, Hassan-Abdi R, Somkhit J, Yanicostas C, Soussi-Yanicostas N. Defective Excitatory/Inhibitory Synaptic Balance and Increased Neuron Apoptosis in a Zebrafish Model of Dravet Syndrome. Cells. 2019;8(10).

34. Tiraboschi E, Martina S, van der Ent W, Grzyb K, Gawel K, Cordero-Maldonado ML, et al. New insights into the early mechanisms of epileptogenesis in a zebrafish model of Dravet syndrome. Epilepsia. 2020;61(3):549–60.

35. Kovac S, Domijan AM, Walker MC, Abramov AY. Prolonged seizure activity impairs mitochondrial bioenergetics and induces cell death. J Cell Sci. 2012;125(Pt 7):1796–806.

36. Kumar MG, Rowley S, Fulton R, Dinday MT, Baraban SC, Patel M. Altered Glycolysis and Mitochondrial Respiration in a Zebrafish Model of Dravet Syndrome. eNeuro. 2016;3(2).

37. Banerji R, Huynh C, Figueroa F, Dinday MT, Baraban SC, Patel M. Enhancing glucose metabolism via gluconeogenesis is therapeutic in a zebrafish model of Dravet syndrome. Brain Commun. 2021;3(1):fcab004.

38. Kim JS, An JY, Lee KS, Chung YA, Choi JS, Lee KH. PET evidence of cerebellar hypometabolism in a patient with familial episodic ataxia-myokymia syndrome. Mov Disord. 2008;23(10):1483–5.

39. D'Adamo MC, Liantonio A, Rolland JF, Pessia M, Imbrici P. Kv1.1 Channelopathies: Pathophysiological Mechanisms and Therapeutic Approaches. Int J Mol Sci. 2020;21(8).

40. Browne DL, Gancher ST, Nutt JG, Brunt ER, Smith EA, Kramer P, et al. Episodic ataxia/myokymia syndrome is associated with point mutations in the human potassium channel gene, KCNA1. Nat Genet. 1994;8(2):136–40.

41. Paulhus K, Ammerman L, Glasscock E. Clinical Spectrum of KCNA1 Mutations: New Insights into Episodic Ataxia and Epilepsy Comorbidity. Int J Mol Sci. 2020;21(8).

42. Aspatwar A, Tolvanen ME, Jokitalo E, Parikka M, Ortutay C, Harjula SK, et al. Abnormal cerebellar development and ataxia in CARP VIII morphant zebrafish. Hum Mol Genet. 2013;22(3):417–32.

43. Trosclair K, Dhaibar HA, Gautier NM, Mishra V, Glasscock E. Neuron-specific Kv1.1 deficiency is sufficient to cause epilepsy, premature death, and cardiorespiratory dysregulation. Neurobiol Dis. 2020;137:104759.

44. Nelson JC, Witze E, Ma Z, Ciocco F, Frerotte A, Randlett O, et al. Acute Regulation of Habituation Learning via Posttranslational Palmitoylation. Curr Biol. 2020;30(14):2729–38.e4.

45. Li D, March ME, Gutierrez-Uzquiza A, Kao C, Seiler C, Pinto E, et al. ARAF recurrent mutation causes central conducting lymphatic anomaly treatable with a MEK inhibitor. Nat Med. 2019;25(7):1116–22.

46. Griffin AL, Jaishankar P, Grandjean JM, Olson SH, Renslo AR, Baraban SC. Zebrafish studies identify serotonin receptors mediating antiepileptic activity in Dravet syndrome. Brain Commun. 2019;1(1):fcz008.

47. MacRae CA, Peterson RT. Zebrafish as tools for drug discovery. Nat Rev Drug Discov. 2015;14(10):721–31.

48. Liu K, Petree C, Requena T, Varshney P, Varshney GK. Expanding the CRISPR Toolbox in Zebrafish for Studying Development and Disease. Front Cell Dev Biol. 2019;7:13.

49. Babin PJ, Goizet C, Raldua D. Zebrafish models of human motor neuron diseases: advantages and limitations. Prog Neurobiol. 2014;118:36–58.

50. Samarasinghe RA, Miranda OA, Buth JE, Mitchell S, Ferando I, Watanabe M, et al. Identification of neural oscillations and epileptiform changes in human brain organoids. Nat Neurosci. 2021;24(10):1488–500.

