## Supplementary Figure 1 (Figure S1) and supplementary figure/movie legends for "*kcna1a* mutant zebrafish as a model of episodic ataxia type 1 and epilepsy"

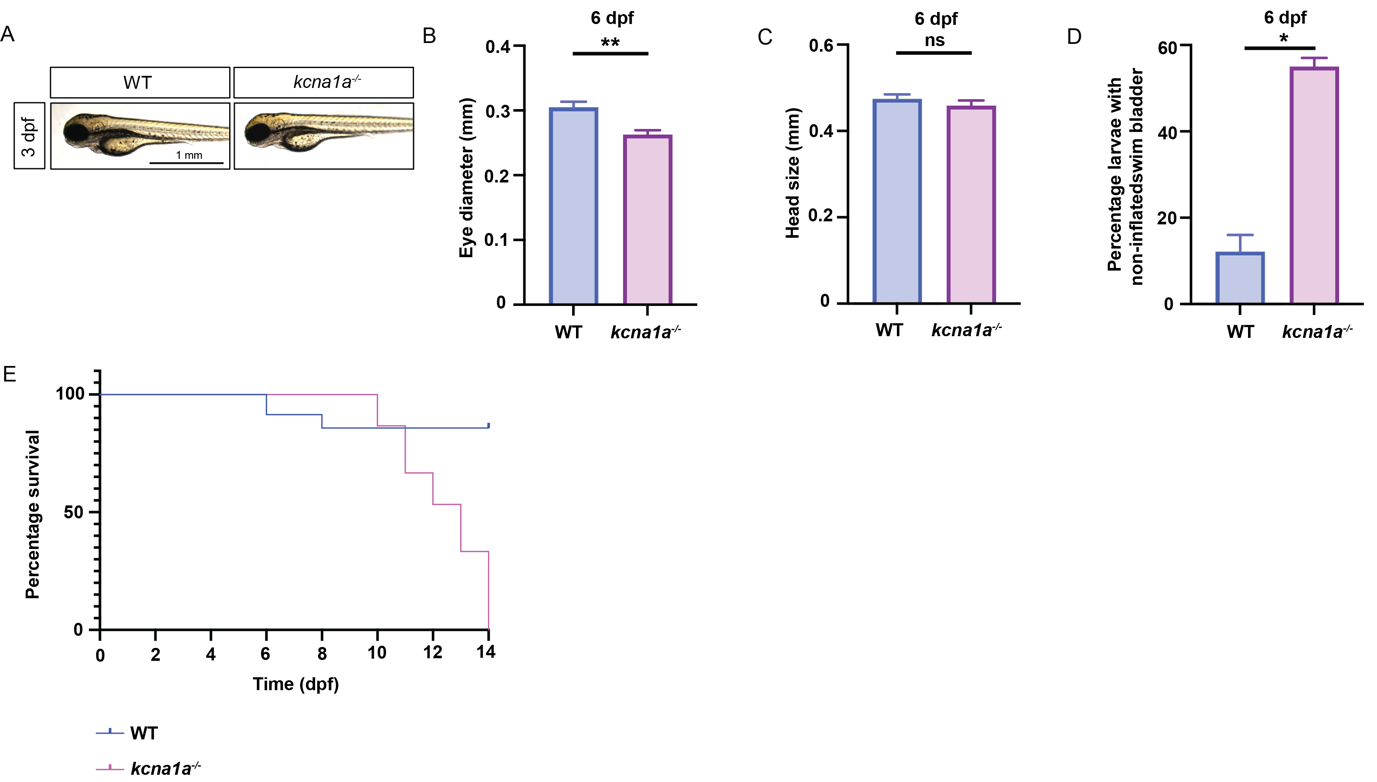

**Supplementary figure 1.** Morphological analysis of *kcna1a^-/-^* and survival curve generation. (A) WT and *kcna1a^-/-^* larvae at 3 dpf. No morphological differences were observed. WT, n=17; *kcna1a^-/-^,* n=26. (B) Quantification of eye diameter in WT and *kcna1a^-/-^* larvae at 6 dpf. *kcna1a^-/-^* show a significant reduction in eye diameter. WT, n=6; *kcna1a^-/-^,* n=10. (C) Quantification of head size in WT and *kcna1a^-/-^* larvae at 6 dpf. No significant differences were observed. WT, n=6; *kcna1a^-/-^,* n=10. (D) Quantification of percentage of WT and *kcna1a^-/-^* larvae at 6 dpf with non-inflated swim bladder. WT, n=270; *kcna1a^-/-^,* n=75. (E) Survival curve of WT and *kcna1a^-/-^*. 100% *kcna1a^-/-^* die by 14 dpf. WT, n=35; *kcna1a^-/-^,* n=15. Data are mean ± s.e.m., ns: no significant changes observed, *P ≤ 0.05, **P ≤ 0.01- Unpaired t-test.

**Movie S1.** 5 dpf WT: Analysis of the pattern of swimming trajectory in 5 dpf WTs. WT larvae exhibit normal swimming pattern at 5 dpf.

**Movie S2.** 5 dpf *kcna1a* mutant: Analysis of the pattern of swimming trajectory in 5 dpf *kcna1a^-/-^*. *kcna1a^-/-^* larvae exhibit abnormal swimming pattern at 5 dpf with uncoordinated movements indicative of ataxia.
